# Discovering lncRNA Mediated Sponge Interactions in Breast Cancer Molecular Subtypes

**DOI:** 10.1101/209015

**Authors:** Gulden Olgun, Ozgur Sahin, Oznur Tastan

## Abstract

**Motivation:** Long non-coding RNAs(lncRNAs) can indirectly regulate mRNAs expression levels by sequestering microRNAs (miRNAs), and act as competing endogenous RNAs (ceRNAs) or as sponges. Previous studies identified lncRNA-mediated sponge interactions in various cancers including the breast cancer. However, breast cancer subtypes are quite distinct in terms of their molecular profiles; therefore, ceRNAs are expected to be subtype-specific as well.

**Results:** To find lncRNA-mediated ceRNA interactions in breast cancer subtypes, we develop an integrative approach. We conduct partial correlation analysis and kernel independence tests on patient gene expression profiles and further refine the candidate interactions with miRNA target information. We find that although there are sponges common to multiple subtypes, there are also distinct subtype-specific interactions. Functional enrichment of mRNAs that participate in these interactions highlights distinct biological processes for different subtypes. Interestingly, some of the ceRNAs also reside in close proximity in the genome; for example, those involving HOX genes, HOTAIR, miR-196a-1 and miR-196a-2. We also discover subtype-specific sponge interactions with high prognostic potential. For instance, when grouping is based on the expression patterns of specific sponge interactions, patients differ significantly in their survival distributions. If on the other hand, patients are grouped based on the individual RNA expression profiles of the sponge participants, they do not exhibit a significant difference in survival. These results can help shed light on subtype-specific mechanisms of breast cancer, and the methodology developed herein can help uncover sponges in other diseases.

## 1 Introduction

Advances in sequencing technologies have revealed that there is a large number of RNAs that do not encode proteins [1]. One class of non-coding RNAs (ncRNAs) comprises microRNAs (miRNAs) that repress gene expression by preferentially binding the compilementary sequence of their target mRNAs [2]. miRNAs play crucial roles in regulating gene expression programs in the normal cell, and their aberrant expression contributes to pathogenesis in several diseases, including cancer. To date, a large number of miRNAs have been shown to be associated with cancer progression, drug resistance or metastasis [3–7].

Another major class of non-coding RNAs is long non-coding RNAs (lncRNAs) that are longer than 200 nucleotides. Although the function of the vast majority of lncRNAs remains to be identified, accumulating evidence suggests that they are highly involved in regulating cellular and pathological processes [8, 9]. Deregulations of several lncRNAs have also been associated with cancer [10, 11]. Recent work has provided evidence for an emerging regulatory role of lncRNAs. By sequestering miRNAs, lncRNAs can reduce the number of miRNAs available for the target mRNA; in this way, they indirectly prevent the target gene repression, acting like a sponge [12, 13]. lncRNA-mediated sponge interactions and their protein-coding targets have been investigated in gastric cancer [15], glioblastoma multiforme [16], pancreatic cancer [17], ovarian cancer [18] and in breast cancer [19].

Identifying sponges by experimental means is a challenge, and the experimental datasets are not available in different contexts such as cancer. Few studies attempt to identify lncRNA sponges computationally. Sponge Scan uses a sequence-based algorithm to detect potential sponge interactions [20]. The algorithm is applicable whenever sequence data is available but does not account for the expression of the RNAs, which provides evidence on the interaction of the RNA species within a given context. Tian et al. [15] find sponge interactions in gastric cancer using microarray expression data. In their approach, they find up-and down-regulated lncRNAs and combine them with miRNA prediction algorithms to construct a lncRNA:miRNA network, but the approach does not consider the correlation structure of RNA expressions. Paci et al. [19] utilizes Pearson correlation and partial correlation analysis to detect sponge interactions in normal and breast cancer samples.

Breast cancer subtypes differ significantly in their molecular profiles and response to therapy. Because miRNAs and mRNAs exhibit different molecular activity patterns in breast cancer subtypes [22, 23, 67], it is expected that there will be subtype-specific lncRNA-mediated sponge interactions as well. Identifying these miRNA sponges can both shed light on the uncharacterized mechanisms of the breast cancer subtypes and potentially help in making better therapeutic decisions. In this work, we use an integrative approach to identify subtype-specific lncRNA:miRNA:mRNA interactions through which lncRNAs compete for binding to shared miRNAs in breast cancer.

To find breast cancer subtype-specific interactions, we systemically analyze lncRNA, miRNA, and mRNA expression profiles of breast cancer patients made available through the Cancer Genome Atlas Project (TCGA) [24]. We first identify statistically related lncRNA:miRNA:mRNA interactions through correlation and partial correlation analysis as in Paci et al. [19] and further refine these candidate interactions using a kernel-based conditional independence test (KCI) [25]. KCI does not assume any parametric form for the random variables that are being tested. Also, for the first time, it is used for finding regulatory interactions. The potential candidate interactions are further filtered in the light of available evidence regarding the miRNA-target interactions. We examine the functional enrichment of mRNAs that participate in sponges, the genomic spatial organization and finally, through the survival analysis of patients, we discover lncRNA-mediated ceRNA interactions with prognostic value.

## 2 Methods

### 2.1 Data Collection and Processing

#### 2.1.1 lncRNA curation

As lncRNAs are not annotated in TCGA, we curated a list of lncRNAs using GENCODE v24 [46]. Based on Gencode24 annotation, 598 of the RNAs present in RNA-Seq expression data are designated as lncRNAs. To minimize erroneous annotations, we further examined each lncRNA’s coding potential with alignment-free method Coding-Potential Assessment Tool (CPAT) [47] and alignment-based method Coding Potential Calculator (CPC) [48]. LncRNAs whose all transcripts are predicted to have high coding potential by both tools are eliminated. The number of lncRNAs that are predicted to have high coding potential by each tool is provided in Figure S1 (Supp. File1).

#### 2.1.2 Expression data processing

Level 3 Illumina HiSeq RNA-seq gene expression and miRNA expression data for human breast cancer were collected from The Cancer Genome Atlas [24] on August 9^th^ 2014 and patient survival data were obtained from the UCSC Cancer Genomics Browser on June 31^st^ 2016. Only the patient data that concurrently include mRNA, lncRNA and miRNA expression data were used. Patients were divided into subtypes based on information in TCGA defined by PAM50. The four subtypes used are Luminal A, Luminal B, Basal, HER2. The number of patients in each subtype is provided in Table S1 (Supp. File1).

In expression data, Reads Per Kilobase Million Reads (RPKM) values were used. To eliminate the genes and miRNAs with very low expression, we assumed that RKPM values below 0.05 are missing and filtered out RNAs that are missing in more than 20 of the samples in each subtype. Expression values are log 2 transformed. RNAs that do not vary across samples were filtered. We eliminated the genes with the median absolute deviation (MAD) below 0.5. MAD is calculated as follows:

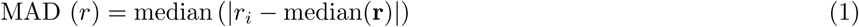

where *r*_*i*_ denotes the RNA expression in sample *i* for RNA *r* and **r** denotes the vector that contains expression values for all samples for RNA *r*.

### 2.2 Statistical Analysis for Finding lncRNA Mediated ceRNA Interactions

To identify ceRNA interactions between lncRNA:miRNA:mRNA, we performed correlation analysis and kernel-based conditional independence test on expression data. Below, random variable *X* denotes the expression level of a lncRNA, random variable *Y* indicates the expression level of a mRNA, and finally random variable *Z* denotes the expression level of a miRNA.

#### 2.2.1 Correlation and Partial Correlation Analysis

For a given ceRNA interaction, we expect expression values of the lncRNA and mRNA to be positively correlated, and if this correlation relies on miRNA expression, the correlation between mRNA, and lncRNA should weaken when miRNA expression is taken into account. To quantify this, first Spearman rank order correlation was calculated between lncRNA and mRNAs, which we denote with *ρ*_*lncRNA, mRNA*_. Next, we calculated the Spearman partial rank order correlation between lncRNA and mRNA, this time controlling for miRNA expression, *ρ*_*lncRNA, mRNA*|*miRNA*_, as follows:

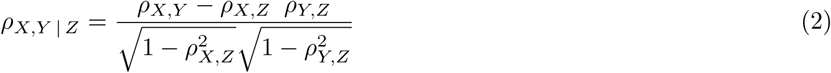

The difference between the correlation and the partial correlation for a miRNA measures the extent the miRNA *Z* is effective in the statistical correlation of *X* and *Y*. This value is calculated:

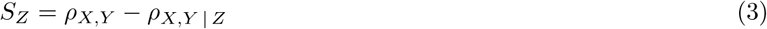

As we look for strongly positively correlated lncRNA and mRNA pairs, only those with correlation *ρ*_*X, Y*_ > 0.5 (*p*-value < 0.05) were considered. Among those, RNA triplets where *S*_*Z*_ is larger than a threshold value, *t*, were retained. We conducted our analysis at two different thresholds *t* =0.2 and *t* =0.3.

#### 2.2.2 Kernel Based Conditional Independence Test

To find lncRNA interactions we also test directly for conditional independence. In a ceRNA interaction, if the interaction of a particular pair of lncRNA (X) and mRNA (Y) were through a shared miRNA (Z), we would expect that lncRNA and mRNA expressions to be conditionally independent given the miRNA expression level. Conditional independence is denoted by *X* _╨_ *Y* | *Z*. Formally, *X* and *Y* are conditionally independent given *Z* if and only if the **P** (*X* | *Y, Z*) = **P** (*X* | *Z*) (or equivalently **P** (*Y* | *X, Z*) = **P** (*Y* | *Z*) or **P** (*X, Y* | *Z*) = **P** (*X* | *Z*) **P** (*Y* | *Z*). That is if *X* and *Y* are conditionally independent given *Z*, further knowing the values of *X* (or *Y*) does not provide any additional evidence about *Y* (or *X*). There are conditional independence tests available for continuous random variables [25, 49–51]. In our work we employ, kernel-based conditional independence (KCI) test proposed by Zhang et al. [25] as it does not make any distributional assumptions on the variables tested. Furthermore, KCI-test does not require explicit estimation of the joint or conditional probability densities and avoids discretization of the continuous random variables, both of which require large sample sizes for an accurate test performance. Below we describe the KCI-test briefly, details of which can be found in [25].

KCI-test defines a test statistic which is calculated from the kernel matrices associated with *X, Y* and *Z* random variables. A kernel function takes two input vectors and returns the dot product of the input vectors in a transformed feature space, *k*: ***χ*** × ***χ*** → ℝ. The feature transformation is denoted by 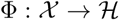 [52], *k*(x_*i*_, x_*j*_) = 〈Φ(x_*i*_) · 〈(x_*j*_)〉 In this work we use the Gaussian kernel, 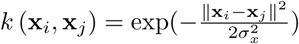, where *σ* > 0 is the kernel width. CI and KCI are based on kernel matrices of *X, Y* and *Z*, which are calculated by evaluating the kernel function for all pairs of samples, i.e. the (i,j)th entry of **K**_*X*_ is *k*(x_*i*_, x_*j*_). The corresponding centralized kernel matrix is 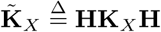 where 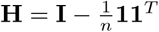 where **I** is the *n* × *n* identity matrix and **1** is a vector 1′s. 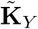 and 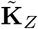 are similarly calculated for *Y* and *Z* variables.

Given the i.i.d. samples **x** ≜ (*x*_1_, *x*_2_, … *x*_*n*_) and **y** ≜ (*y*_1_, *y*_2_, … *y*_*n*_) the unconditional kernel test first calculates the centralized kernel matrices, 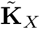 and 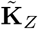 from the samples **x** and **y** and then eigenvalues of the centralized matrices. The eigenvalue decompositions of centralized kernel matrices 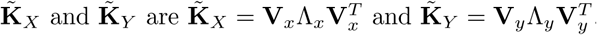 Here Λ_*x*_ and Λ_*y*_ are the diagonal matrices containing the non-negative eigenvalues λ_x, *i*_ and λ_*y, j*_ in descending order, respectively. **V**_*x*_ and **V**_*y*_ matrices contain the corresponding eigenvectors. Zhang et al. [25] show that under the null hypothesis that X and Y are independent, the following test statistic:

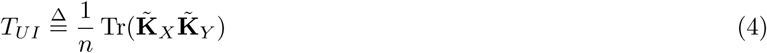

has the same asymptotic distribution (*n* → ∞) as

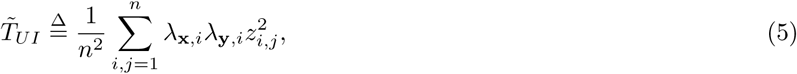

Here *z*_*i, j*_ are i.i.d. standard Gaussian variables, thus 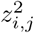 are i.i.d 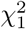 - distributed. The unconditional independence test procedure involves calculating *T*_*UI*_ according to Eq (4). Empirical null distribution of 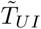 is simulated by drawing i.i.d random samples for 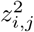 variable from 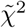. Finally, the *p*-value is calculated by locating *T*_*UI*_ in the empirical null distribution.

The kernel conditional independence test also makes use of the centralized kernel matrices. Under the null hypothesis that *X* and *Y* are conditionally independent given *Z*, the following test statistic is calculated:

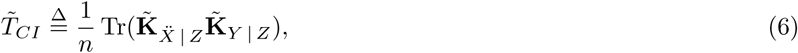

where 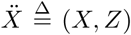 and 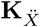 is the centralized kernel matrix for 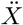. As Zhang et al. [25] report has the same asymptotic distribution as

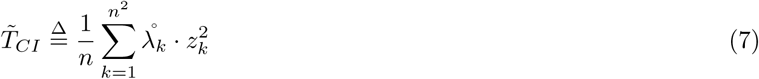

The details of the definition of 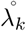 and 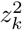 can be found in [25]. The procedure involves calculating the empirical *p*-value based on the test statistic as defined in Eq (4) and simulating the null distribution based on Eq (7).

Using the unconditional kernel independence test, we first test the null hypothesis that a lncRNA and mRNA pair is independent. We consider pairs for whom the null hypothesis is rejected at the 0.01 significance level. For each of the lncRNA:mRNA pair, we test their conditional independence with all miRNAs. A lncRNA and mRNA pair is tested for conditional independence given a miRNA using KCI. Those pairs that are found to be independent at significance level 0.01 are considered as potential lncRNA-mediated ceRNAs.

### 2.3 Filtering ceRNAs Based on miRNA-Target Interactions

To identify interactions that are biologically meaningful, we filtered the potential ceRNA interactions that were not supported by miRNA target information. The miRNA:mRNA and miRNA:lncRNA interactions are retrieved from multiple databases as listed in Table 1. The candidate sponges are retained if both mRNA and lncRNA have support for being targeted by the miRNA of the sponge.

**Table 1:**
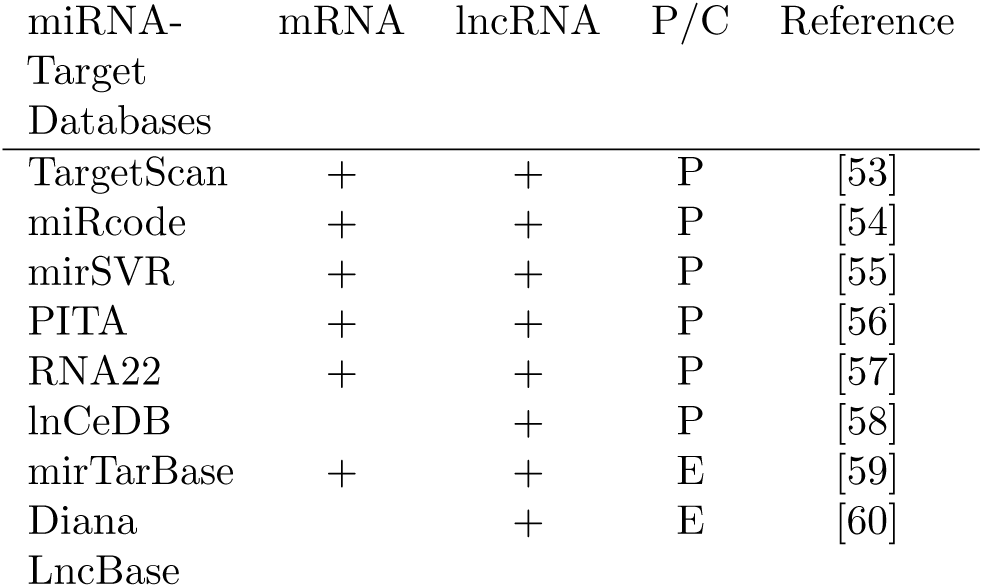
Computationally and experimentally validated miRNA-target databases used for mRNA and lncRNA. Plus signs denote databases that are used for the miRNA interactions of the RNA type. ‘P’ denotes predicted target information while ‘E’ denotes experimentally supported target information.

### 2.4 Identifying ceRNAs with Prognostic Value

To evaluate the ceRNA interactions in terms of their prognostic potential, we analyzed the survival of the patients based on the expression patterns of each sponge interaction. In a sponge interaction, we expect the lncRNA and mRNA to be regulated in the same direction and miRNA to be in the opposite direction. For each ceRNA found in a subtype, the patients are divided into two groups based on the regulation patterns of the RNAs that participate in the ceRNA. For the up-down-up pattern, the first group comprises patients whose sponge lncRNA and mRNA are up-regulated, and miRNA is down-regulated; the second group includes all patients that do not fit in this pattern. Similarly, we divide the patients based on the down-up-down pattern: if both lncRNA and mRNA are down-regulated whereas miRNA is unregulated, such patients constitute one group, and the rest of the patients constitute the second group.

For each of the subtype-specific ceRNAs identified, we tested whether the patient groups’ survival rates are significantly different from each other using log-rank test [61] (*p*-value < 0.05). We further excluded ceRNAs if any of the RNAs can by itself divide the patients into groups that differ in terms of their survival distributions significantly. In each subtype, we split the patients as up-regulated and down-regulated for each of the RNA participating in the ceRNA interaction separately. If at least one of the molecules leads to groups with significant survival difference (log-rank test, (*p*-value < 0.05)), we disregard this ceRNA from the list of prognostic ceRNAs. This last step ensures that the prognostic difference is due to the interactions between the RNAs but not stem from the expression of the single RNA’s expression patterns.

The identified ceRNA interactions are further divided based on the following *ƒ* score that reflects the interaction’s prognostic value with respect to the single RNA’s prognostic value:

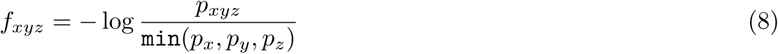

Here, *p*_*xyz*_ is the *p*-value attained in testing whether patient survivals differ based on the log-rank test whereas *p*_*x*_, *p*_*y*_, *p*_*z*_ indicate the *p*-values obtained by testing patient survival distribution differences due to lncRNA, mRNA, and miRNA expression patterns, respectively.

In the above analysis, RNAs that have expression levels above (or below) a particular threshold value are considered up-regulated (or down-regulated). This threshold value is selected among the candidate cut-off values of expression as the one that results in the lowest *p*-value in the log-rank test when patients are divided based on this cut-off. The candidate cut-off values are the 10^th^ and 90^th^ percentiles, mean, median or the lower and upper quartiles of the expression values of the patients in each subtype.

### 2.5 Pathway and GO Enrichment Analysis

We conducted pathway and GO enrichment of mRNAs that participate in subtype-specific sponges. Enrichment tests are conducted with clusterProfiler [62] with Bonferroni multiple hypothesis test correction. In deciding enriched pathways and GO terms, a *p*-value cut-off of 0.05 and FDR cut-off of 1 × 10^−4^ are used. In both pathway and GO enrichment analysis the background genes were the union of mRNAs that remained after MAD filtering step (Step B in Figure 1(A)). For pathway enrichment analysis, different pathway data sources were downloaded from Baderlab GeneSets Collection [63]. List of all pathways that are employed in this analysis is provided in Table S2 (Supp. File1). Redundant pathways are eliminated when different sources are combined. Additionally, a pathway enrichment analysis is conducted with KEGG pathways (downloaded on February 28^th^ 2017).

**Figure 1:**
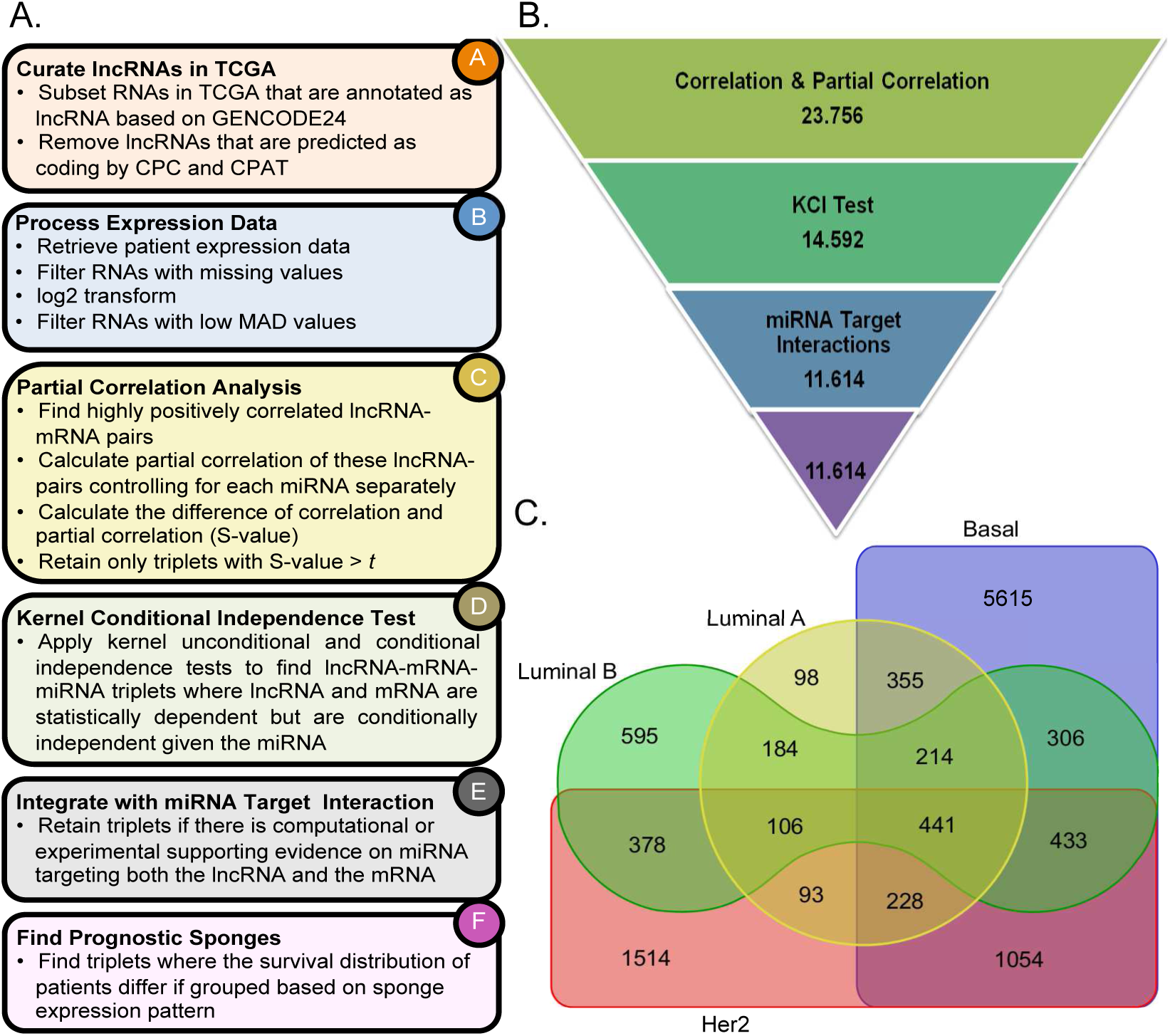
A)Overview of the methodology, each box represents a step in the methodology. Steps B-F are conducted for each breast cancer subtype separately. B)The number of ceRNAs remained after each main filtering step when *t* = 0.2 (Step C in Figure 1A). C)Venn diagram of ceRNA interactions discovered in each of the breast cancer molecular subtype.

### 2.6 Clustering mRNAs

If mRNAs are highly correlated with each other, we often find that correlated mRNAs participate in ceRNA interactions with the lncRNA and miRNA pair. We consider the mRNAs that participate in a ceRNA interaction with the same pair of lncRNA and miRNA. If all mRNAs are strongly correlated among each other, where all the pairwise correlations are above than 0.7, all mRNAs are assigned into the same cluster. Otherwise, we apply Ward hierarchical clustering method to find groups of correlated mRNAs [64]. We determine the optimal number of clusters with Mojena’s stopping rule [65] using Milligan and Cooper’s [66] correction.

## 3 Results

### 3.1 Overview of Discovered ceRNA Interactions

To discover subtype-specific breast cancer ceRNA interactions, we employ the methodology summarized in Figure 1(A) and identify ceRNAs specific to four molecular subtypes of breast cancer: Luminal A, Luminal B, HER2, and Basal. The number of candidate ceRNA interactions that remain after each main step when in the partial correlation analysis step *S* value threshold *t* = 0.2 is employed, is provided in Figure 1(B) (see Figure S2(A) in Supp. File1 for *t* = 0.3). The total number of ceRNA interactions found in all subtypes is 11.614. Figure 1(C) shows the Venn diagram of number of ceRNA interactions discovered for the four subtypes (see Figure S2(B) in Supp. File1 for *t* = 0.3).

Although there are sponges that are detected in multiple subtypes, there are also a large number of sponges that are only specific to a single subtype (Table S3 and Figures S3A and S3B in Supp. File1). The list of sponges identified in each subtype, their partial correlation analysis, KCI-test results and target information are provided in Supp. File 2.

We analyze the specificity of the individual RNAs that participate in each of the subtypes. Figures 2A and 2B display the number of sponges per lncRNA and miRNA for *t* = 0.2 (Figure S3C in Supp. File1 for *t* = 0.3). Some lncRNAs and miRNAs participate in sponges of all the subtypes (Table 2); i.e., KIAA0125 participates in a large number of sponges across the four subtypes. KIAA0125 has been reported to act as an oncogene in bladder cancer related to cell migration and invasion [26]; however, no functional relevance to breast cancer has been reported to date. HOTAIR, which is one of the lncRNAs that has been associated with metastasis [27], is found to participate in sponges of all the subtypes except HER2. Similarly, miRNAs hsa-miR-142, hsa-miR-150, and hsa-miR-155 participate in ceRNA interactions of all subtypes. There are also RNAs that take part in sponges exclusively in a single subtype (Table S4 in Supp. File1). For example, the lncRNA C17orf44 is specific to HER2 (Figure 2(A)) while hsa-miR-342 is only found in Basal ceRNA interactions (Figure 2(B)). Several studies indicated that miR-342 is linked to BRCA1 mutated breast cancer, most of which are the Basal subtype [28–30]. Similarly, some mRNAs are involved in ceRNA interactions only in a single subtype (see Supp. File 3 for all the mRNAs in the interactions and see for only the prognostic mRNAs see Supp. File 4). These subtype-specific RNAs are of great value for understanding the dysregulated cellular mechanisms in each subtype.

**Figure 2:**
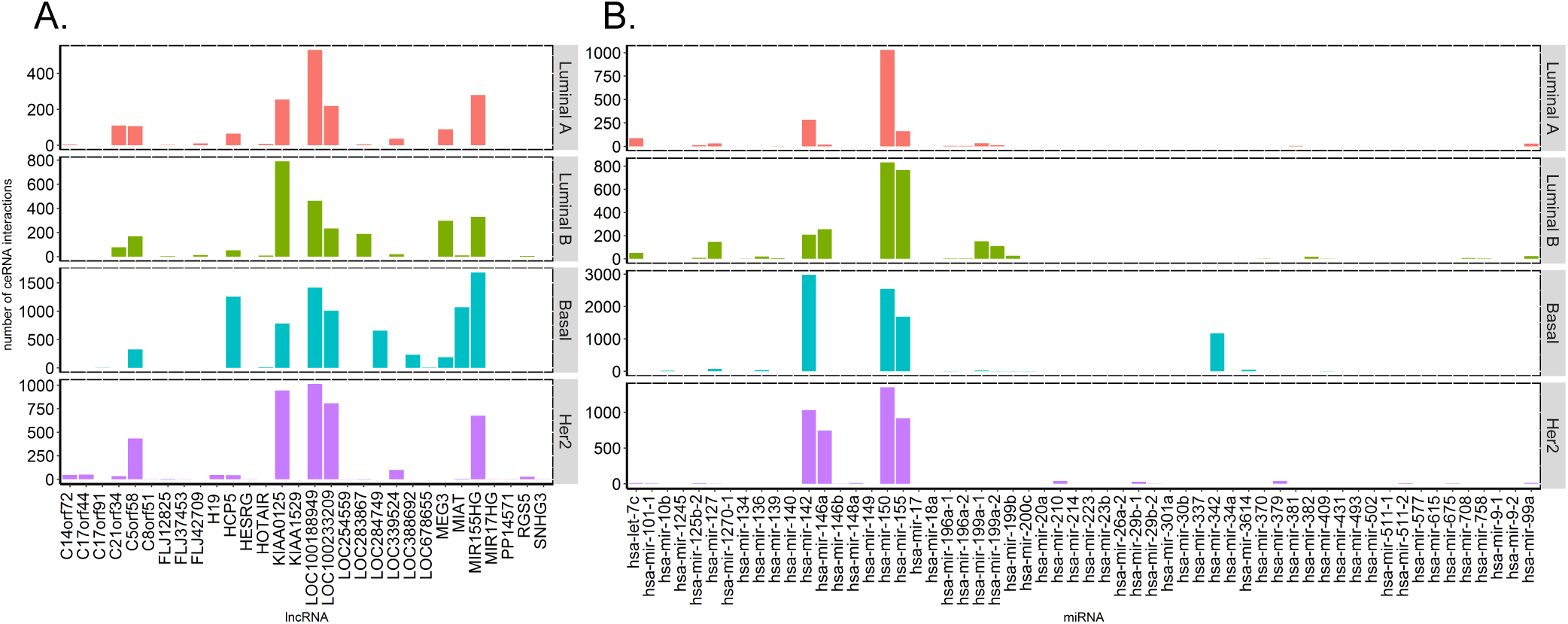
Number of ceRNA interactions discovered that A) lncRNAs and B) miRNAs take part in each breast cancer subtype (*t* = 0.2).

**Table 2:** List of lncRNAs & miRNAs that are found to participate in sponges of all four subtypes.

| miRNA | lncRNA |
| --- | --- |
| hsa-miR-142 | LOC100188949 |
| hsa-miR-196a-1 | C5orf58 |
| hsa-miR-127 | LOC100233209 |
| hsa-miR-155 | HCP5 |
| hsa-miR-150 | KIAA0125 |
| hsa-miR-196a-2 | C21orf34 |
| hsa-miR-125b-2 | MIR155HG |

The lncRNA:mRNA networks for each subtype are shown in Figure S4 (Supp. File1) and Supp. File 8. In these networks, each node denotes a lncRNA or an mRNA while an edge represents an interaction through a shared miRNA. The number of nodes and edges are provided in Table S5 (Supp. File1). In Luminal A, lncRNA LOC100188949 regulates the majority of the sponge interactions, while C21orf34 also form a smaller connected component of its own. In Luminal B, KIAA0125 is at the center of the many interactions while a few other lncRNAs among them are HOTAIR and C21orf34 mediates a small number of interactions. Basal and HER2 subtypes include a large number of interactions. In Basal subtype, among others HCP5, MIR155HG, MIAT are the hubs of the network. In HER2 subtype KIAA0125, LOC100188949 and LOC100233209 are the top 3 largest hubs.

We find that the same lncRNA:miRNA pair participates in multiple sponge interaction with different mRNAs. As an example, HER2 subtype-specific C14orf72-hsa-miR-150 lncRNA:miRNA pair interacts with 45 different mRNAs, the same is not true for lncRNA:mRNA pairs. The number of ceRNA interactions per lncRNA:miRNA pairs is provided in Figure S5 (Supp. File1). We also analyze the data by clustering mRNAs that participate in a sponge with the same lncRNA:miRNA pair based on mRNA expression correlation. The list of the identified sponges in the view of these clusters are provided in Supp. File 5.

### 3.2 Spatially Proximal ceRNAs Interactions

Although regulatory interactions can take place between molecules encoded in different chromosomes, spatial proximity often hints a tight regulatory coordination. We examine the genomic locations of the RNAs that participate in a sponge interaction. Those sponges for which the participating RNAs are within the 100KB distance of each other are identified. The most striking case is the set of sponge interactions that take place between HOTAIR, hsa-miR-196a miRNAs and HOXC genes (Figure 3(A)). These sponge interactions are discovered in all subtypes except HER2. Both HOTAIR, has-miR-196a-1, and hsa-miR-196a-2 are all spatially proximal to the HOX gene clusters on chromosome 12 (Figure 3(B)). HOX genes are highly conserved transcription factors that take master regulatory roles in numerous cellular processes including development, apoptosis, receptor signaling, differentiation, motility, and angiogenesis. Their aberrant expression has been reported in multiple cancer types [31]. HOXA is reported to have altered expression in breast and ovarian cancers; other HOX genes are also associated with multiple tumor types, including colon, lung, and prostate cancer. The lncRNA partner of this sponge interaction is HOTAIR. Up-regulation of HOTAIR is associated with metastatic progression and low survival rates in breast, colon, and liver cancer patients [13–16, 18, 21, 26, 35, 36]. The complete list of sponge interactions whose members exhibit such spatial proximity at least between two RNAs in the sponge is provided in Supp. File 6.

**Figure 3:**
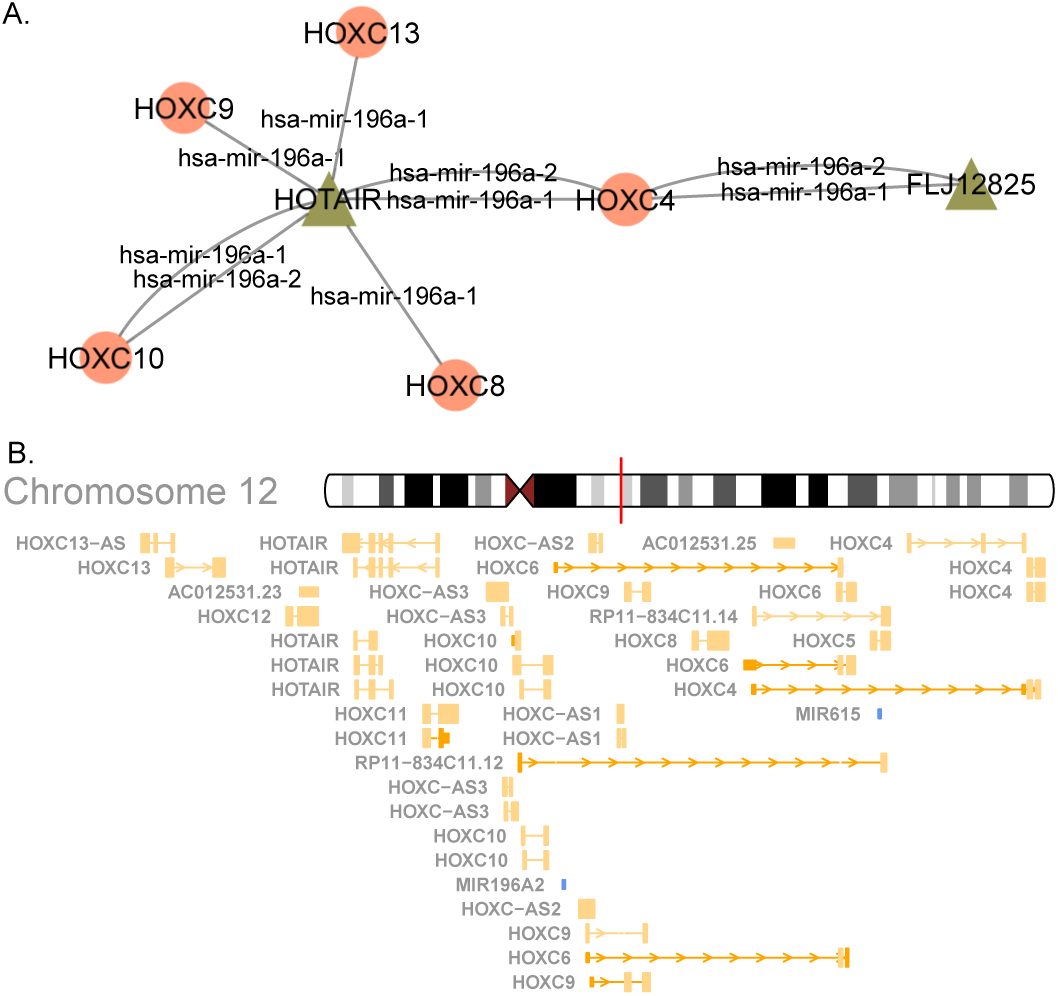
A) The network of sponge interactions between HOTAIR, hsa-miR-196a miRNAs and HOXC genes. The circles denote lncRNAs, and triangles denote mRNAs. An edge exists between a lncRNA and an mRNA if there is a sponge interaction between them; the edge label indicates the miRNA that regulates the interaction. B) The genomic locations of the sponge interactions on chromosome 12.

### 3.3 Functional Enrichment Analysis of mRNAs in ceRNAs

To understand the patterns of pathways related to identified sponges, we conducted pathway enrichment of mRNAs that participate in the sponges separately. The top enriched pathways are found to be common across subtypes (see S6-S7 Figures, Supp. File1) and these pathways are mostly related to the immune system and signaling pathways, which are essential modulators of cancer progression and therapy response [37]. Interestingly, interferon alpha/beta signaling pathway is among the top pathways for Basal subtype (*p*-value 7.20 × 10^−23^) while it is not found enriched in other subtypes (*p*-value cut-off 0.05 and FDR cutoff 1 × 10^−4^). Considering the key role of interferon signaling in the immune system, and the positive correlation between immune cell infiltration and aggressiveness of Basal subtype of breast cancer, our results suggest that mRNAs involved in ceRNA interactions might contribute to the different immune profile of the Basal subtype [38, 39]. Complement cascade induces cell proliferation which causes carcinogenesis including invasion, cell death, and metastasis [40], which are Basal subtype characteristics. We detected C2, C3, C3AR1, C4A, C7 complement genes in Basal ceRNA interactions. Consequently, complement cascade pathway may be significant for the Basal subtype.

The overlap between the enriched pathways in different subtypes is shown on a Venn diagram (Figure S8 in Supp. File1). The list of pathways that are found enriched only in a single subtype is listed in Table S7 in Supp. File1 with p-value cut-off 0.05 and FDR cutoff 1 × 10^−4^. Interestingly, the PI3K pathway is found to be enriched specifically in Luminal A. This is interesting as the most frequently mutated gene in Luminal A is PIK3CA (45% of the patients in TCGA), and there are PIK3CA mutations that are specific to this subtype [24]. This suggests that ceRNA interactions might be key regulators of the PI3K pathway, especially in this subtype of tumors which comprises of more than 60% of breast cancers.

Integrin signaling is widely studied in breast cancer literature since integrins incorporate breast cancer progression [42]. Moreover, integrins play key roles in migration, invasion, and metastasis of cancer cells. Enrichment of integring signaling in mRNAs involved in ceRNA networks might suggest that HER2-specific ceRNA interactions might contribute the aggressive progression of HER2 subtype, similar to the Basal subtype of breast cancer. Thus, they drive tumor cell to metastasis [42]. HER2 subtype-specific enriched pathways contain integrin signaling pathways (TableS7 in Supp. File1).

### 3.4 Prognostic Sponge Interactions

To identify ceRNA interactions with prognostic value, for each of the identified sponge we checked whether the sponge expression pattern divides the patients into groups that differ in their survival probability. To further verify that this difference is due to the interaction and not due to an individual RNA molecule that participates in the sponge, we set another filter. We only consider interactions where there is a significant difference in survival when patients are grouped based on ceRNA expression pattern while there is no significance due to a grouping based on a single RNA molecules’ expression pattern. These prognostic ceRNA interactions are ranked based on *ƒ*-score (details in 2.4) and are provided as a list in Supp. File 7 and the network of interactions are shown in Figure 10 in Supp. File1. KM plots for the two examples are shown in Figure 4. The distribution of lncRNAs that participate in the prognostic sponges are provided in Figure S9 in Supp File1, and the network of interaction among prognostic sponges are provided in Figure 5 and the Supp. Cytoscape file in Supp. File 9. For example, patients with a sponge pattern where MEG3 and COL12A1 are high and miR-1245 low have better survival than other patients. Furthermore, none of these three RNA molecules is able to separate the patients into groups that differ in survival probabilities. This suggests that examining key sponge patterns might have a better prognostic value than that of individual genes.

**Figure 4:**
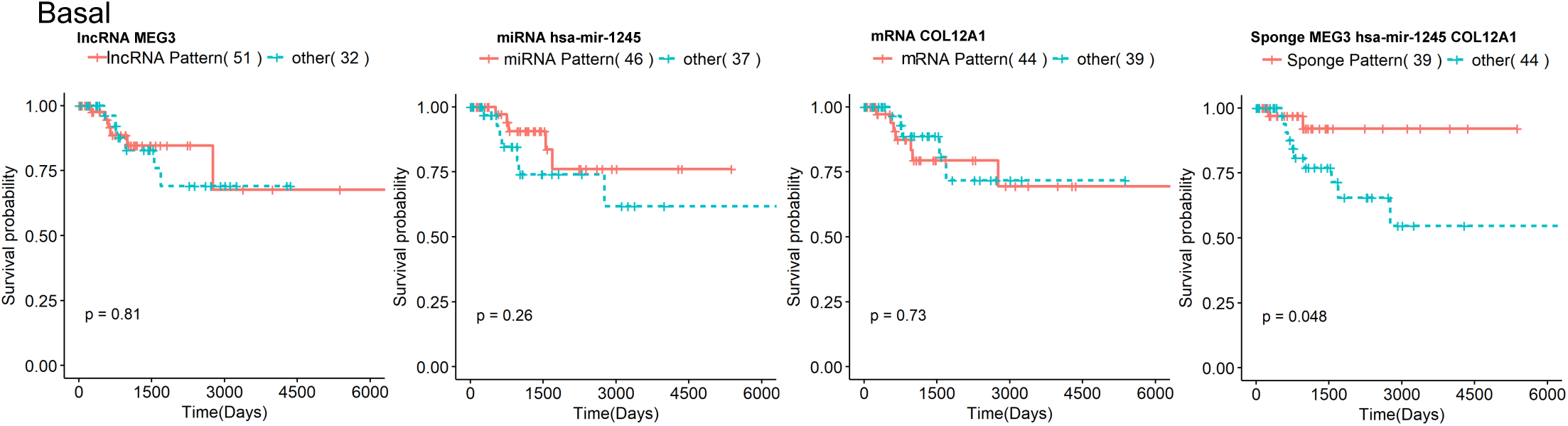
Kaplan Meir survival plots when patients are divided based on individual expression patterns of the RNAs (the first three plots in each panel) and when patients are divided based on the sponge expression pattern (4th plot) for MEG3, hsa-miR-1245, COL12A1 sponge

**Figure 5:**
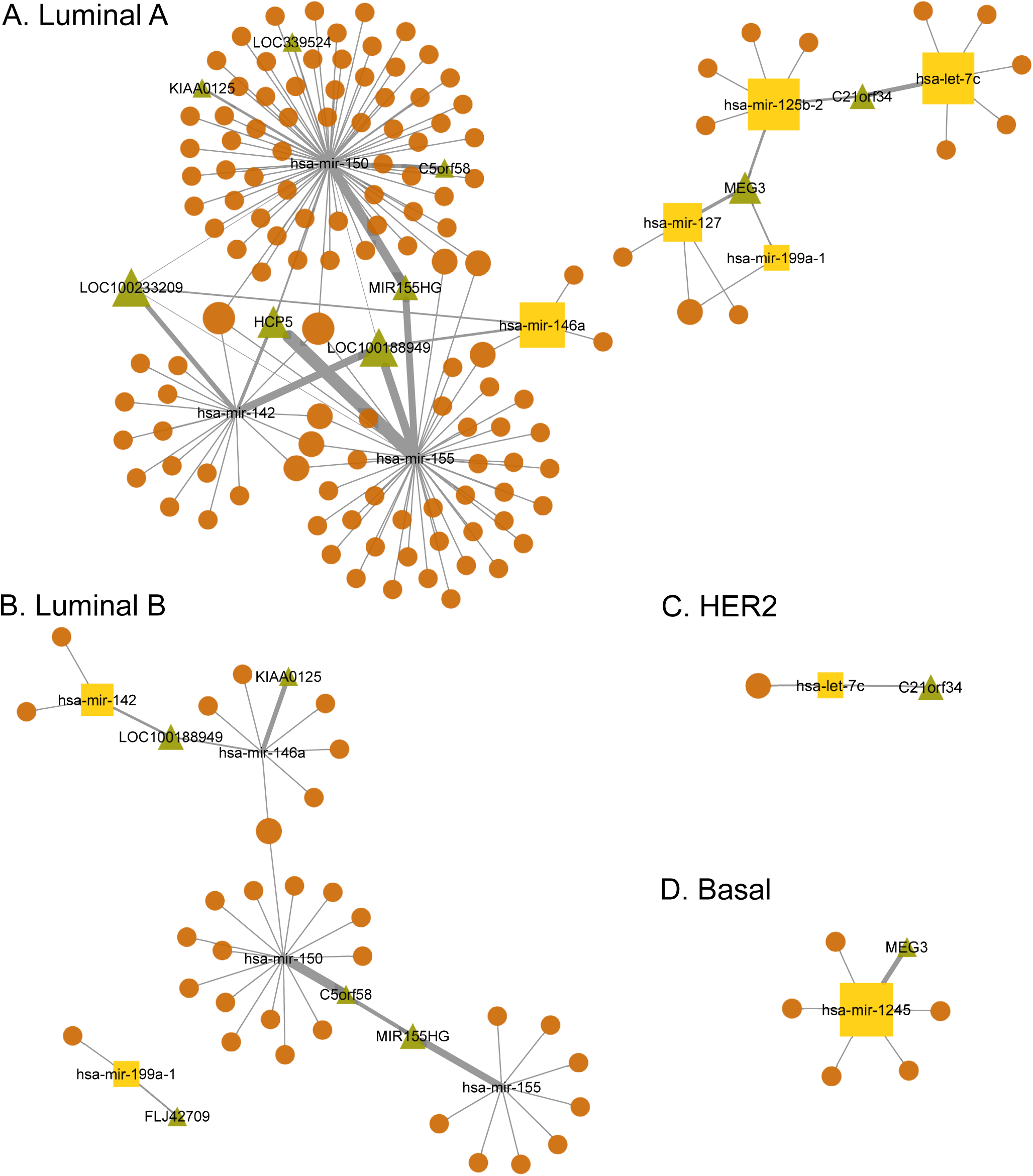
lncRNA:miRNA:mRNA network for all breast cancer subtypes. lncRNAs are represented by the green triangle symbol, mRNAs are represented by orange ellipse symbol and miRNAs are with the yellow rectangle. Each node size is scaled by its degree, the number of edges incident to the nodes and edge width is scaled by the number of occurrence of the node pair. The network was constructed using the Cytoscape (v3.4.0) [45].

## 4 Discussion and Conclusion

As transcriptome is cataloged with greater depth, it has become evident that the vast majority of the mammalian transcriptome is non-coding. One type of non-coding RNAs is lncRNA. A growing body of evidence demonstrates that lncRNAs are deregulated in cancer just like mRNAs and miRNAs [43]. For example, over-expression of the lncRNA HOTAIR in breast cancer patients is reported to be highly predictive of patient survival and progression to metastasis [27]. An emerging role of lncRNAs is that they compete for binding to miRNAs, acting as a sponge to regulate the gene activity. This three-way regulatory interaction between lncRNAs, miRNAs, and the mRNAs is observed in multiple cancer types, including breast cancer. Our contribution is in this work is two folds. Firstly, we identify potential subtype-specific lncRNA mediated sponge interactions in breast cancer. Secondly, we develop an integrative methodology, which can also be applied to other diseases.

Our method uses rigorous statistical tests on patient RNA expression profiles to find the statistical relationship required among the triplets of lncRNA, mRNA, and miRNAs. This candidate list is compiled by using the miRNA target information. To this end, we retrieve experimentally validated and computationally predicted miRNA:lncRNA and miRNA:mRNA information from multiple databases. Due to the limited knowledge on experimentally confirmed interactions, we choose to include predicted targets in our analysis, as well. As more experimental target information becomes available, the identified list can be further filtered to reduce the false positive rate. To assist further analysis, we also curate additional information such as the number of databases supporting a predicted interaction and the number of miRNA binding sites in the lncRNA partner. This additional information can also be used to subdivide the lists further into subsets.

Experimentally validated and lncRNA mediated sponge interactions for breast cancer subtypes are not readily available. This absence limits the efforts to assess the statistical power and false positive rate of our method and complicates the choice of cut-off values used in the compilation of the final candidate list. For this reason, we report our results at two cut-off values that differ in their stringency. We also perform several analyses to investigate the relevance of the discovered potential interactions and to validate the interaction list with indirect supporting evidence. Firstly, the functional enrichment analysis of mRNAs that involve subtype-specific sponges are conducted. This analysis reveals subtype specific mRNA partners are enriched with pathways/processes known to be specific to some of the subtypes. We consider this as an indirect validation. Secondly, the spatial organization of the RNA triplets that participate in the genome reveals that some of the sponges are positioned in close proximity of each other on the genome, hinting a regulatory relationship between these RNAs. Thirdly, a subset of the interactions is found to have prognostic value. Based on the sponge expression patterns, patients can be divided into two groups that differ in terms of their survivals.

These findings can be probed and tested by experimental analyses and potentially help uncover unknown molecular mechanisms of breast cancer subtypes. The methodology developed herein can also be applied to other diseases and cellular states.

